# What is the state space of the world for real animals?

**DOI:** 10.1101/2021.02.07.430001

**Authors:** Vijay Mohan K Namboodiri

## Abstract

A key concept in reinforcement learning (RL) is that of a state space. A state space is an abstract representation of the world using which statistical relations in the world can be described. The simplest form of RL, model free RL, is widely applied to explain animal behavior in numerous neuroscientific studies. More complex RL versions assume that animals build and store an explicit model of the world in memory. To apply these approaches to explain animal behavior, typical neuroscientific RL models make assumptions about the underlying state space formed by animals, especially regarding the representation of time. Here, we explicitly list these assumptions and show that they have several problematic implications. We propose a solution for these problems by using a continuous time Markov renewal process model of the state space. We hope that our explicit treatment results in a serious consideration of these issues when applying RL models to real animals.

---

Predicting rewards is essential for the sustained fitness of animals. Since animals (including humans) experience events in their life along the continuously flowing dimension of time, predicting rewards fundamentally requires a consideration of this timeline (**Fig 1**). Several models have been proposed for how animals learn to predict rewards based on their experience (Balsam et al., 2010; Brandon et al., 2003; Dayan, 1993; Gallistel and Gibbon, 2000; Gallistel et al., 2019; Pearce and Hall, 1980; Rescorla and Wagner, 1972; Schultz, 2016; Sutton and Barto, 1998; Wagner, 1981). Among these, the most widely used class of models in neuroscience is reinforcement learning (RL). The core concept of RL models is that animals make an initial prediction about upcoming reward, calculate a prediction error — the difference between the experienced reward and the predicted reward, and then update their prediction based on this prediction error. While RL models were inspired by psychological models such as the Rescorla-Wagner (Rescorla and Wagner, 1972) or the Pearce-Hall (Pearce and Hall, 1980) models, mathematically rigorous versions of it have borrowed extensively from concepts in computer science. Briefly, RL contends that the learning agent (e.g. real animals) represents the structure of their world in a “state space” that abides by simplified principles such as Markov chains (Niv, 2009; Sutton and Barto, 1998). A state is any abstract representation of observable or unobservable events in their world, and Markov chains are a special kind of state space in which transitions between states do not depend on the history of previously experienced states. Commonly used Markov chain models discretize the flow of time (Niv, 2009; Schultz et al., 1997) or assume temporal basis functions during intervals between events (Gershman et al., 2014; Ludvig et al., 2008, 2012; Petter et al., 2018). These formulations have an intrinsic mathematical simplicity to them, which makes rigorous mathematical calculations possible (e.g. the Bellman equation for value update). Here, we show that these simplifying assumptions have problematic implications when applied to learning in real animals, as they often do not naturally account for the timeline of experience of real animals. Our hope is that the explicit treatment considered here stimulates serious considerations of these issues. As a possible solution to these issues, we present an alternate formulation of the state space for animals that provides a rich representation of their world, while naturally accounting for the continuous passage of time.

**Fig 1.**
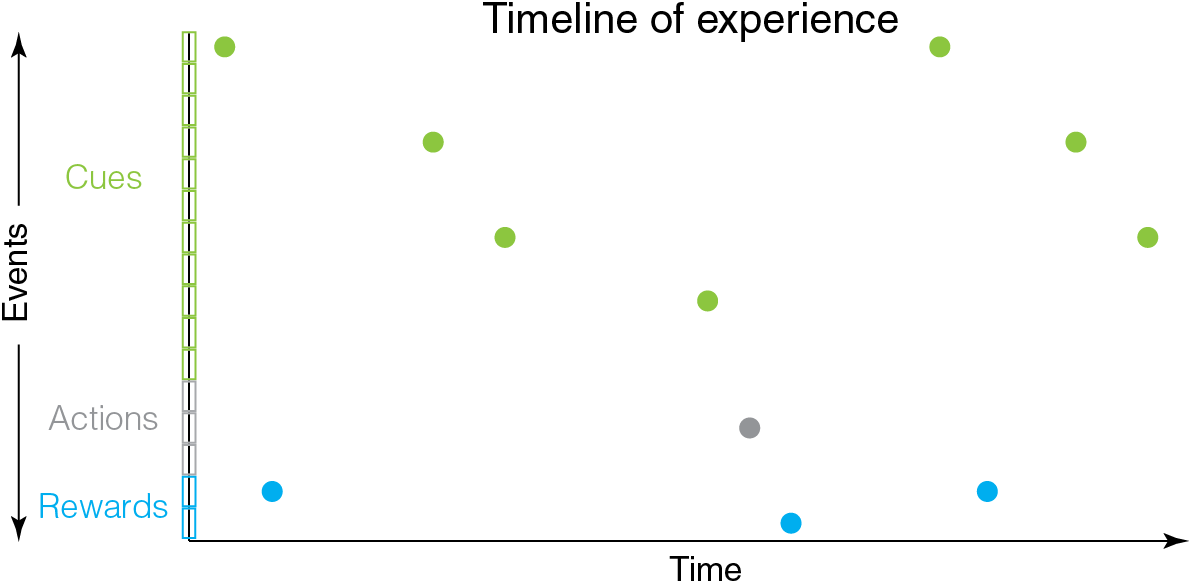
Animals experience events in their life in a timeline along the continuously flowing dimension of time. Thus, prediction of rewards requires a consideration of the flow of time. Here, external cues, internally generated actions and rewards are shown by separate colors. Distinct types of events within these groups are shown by individual boxes along the y-axis.

## Example illustrative task

Perhaps the simplest RL task for animals is cue-reward learning. Most commonly, this is studied in Pavlovian conditioning experiments in which an environmental cue is predictive of an upcoming reward (Pavlov, 1927). Often, there is a delay between when the cue turns off and the subsequent reward delivery (e.g. Bangasser et al., 2006; Beylin et al., 2001; Coddington and Dudman, 2018; Kobayashi and Schultz, 2008; Schultz et al., 1997). This variant of the task is known as trace conditioning. We will use this simple illustrative example throughout this paper. The main reason for doing so is to show that even the simplest tasks require problematic assumptions. Indeed, the problems laid out here become more severe for tasks requiring reward predictions based on actions. Another reason is that this type of learning, i.e. cue followed by delay followed by reward, is highly ethologically relevant. For instance, for wild foragers, environmental landmarks can often act as “cues” predictive of a reward after some distance (or delay) (Chittka et al., 1995; Wystrach et al., 2019a, 2019b). Similarly, for many animals, cues reflecting the end of winter are predictive of an increased availability of food reward. It is then perhaps not surprising that even insects show evidence of such learning (Chittka et al., 1995; Dylla et al., 2013; Menzel, 2012; Toure et al., 2020; Wystrach et al., 2019a). We will first discuss the common mathematical formulation for representing state space in this task, before discussing implicit assumptions and their problematic implications.

## Markov Chains

The mathematical concept of Markov chains is the building block for state space representations in RL. Briefly, a Markov chain is formed by a set of states, *S* = *{1*, *2*,…, *n}*. An implicit assumption is that all the relevant states in the world have been specified in *S*. The process is assumed to start from one state and successively moves to another state (possibly itself) with a probability *p*_*ij*_ (where i and j are indices for the starting and ending states and can be equal). Each move is called a step. Each step results in a transition, which could be a self-transition to the same state. Crucially, the “transition probabilities” from a given state do not depend on the history of states. If we index the step number (a measure of time) by a subscript *t*, this means that *p*(*s*_*t*_|*s*_*t-1*_, *s*_*t-2*_,…, *s*_*1*_)=*p*(*s*_*t*_|*s*_*t-1*_). This absence of history dependence is known as the Markov property, and allows some convenient mathematical representations.

The state space in RL is typically such a Markov chain. In more realistic RL formulations, the animal can also take a set of actions *A*=*{a*_*1*_,…,*a*_*m*_*}* that transitions the agent from one state to another with a conditional probability of *p(s*_*j*_ | *s*_*i*_, *a*_*k*_). These transition probabilities can collectively be represented by a transition matrix *P*. The state space for an RL agent is fully described by *S, A and P*. This more general state space that includes an ability of agents to interact with its states using actions is the Markov Decision Process used in RL.

For simplicity, we will only consider the example illustrative task discussed above, in which a reward follows a cue after a delay. Hence, we will omit considerations of actions and the dependence of *P* on actions. In this task, the states can be minimally specified as the cue state and the reward state. Representing these stimuli as states allow an animal to store the sensory properties of these states in memory. For instance, the animal could learn that an auditory cue has a specific set of sensory attributes such as frequency profile, loudness, duration etc. Similarly, the sensory properties of a type of reward can be represented as a reward state. These various attributes can be stored as part of the memory of that state. Additionally, it is assumed that animals learn a representation of a scalar value for reward. In RL, the reward values are typically denoted by *R(s, a),* a scalar value associated with each state-action pair. For our purpose, we will denote the reward function by *R(s)*. For the cue and reward state formalism that we adopt, *R(cue)=0* and *R(reward)=reward value*. Thus, *S, P, and R* completely describe the cue-reward task of interest.

## Dealing with time in Markov chains

The biggest problem with the above state space model is that there is no representation of time. The task of interest contains a delay between the cue and reward, and a delay from the reward to the next presentation of the cue (typically called the intertrial interval or ITI). However, there is no representation of these delays in the above Markov chain.

Neuroscience-related RL models solve this problem using the idea of “microstates” (**Fig 2**). The simplest such model assumes that the delay from cue to reward is represented by a series of states of equal duration (example 1 in **Fig 2**). This is known as the complete serial compound model of the state space (Moore et al., 1998; Schultz et al., 1997; Sutton and Barto, 1990). Here, the set of states is *S={cue, delay*^*1*^, *delay*^*2*^,… ,*delay*^*n*^, *reward}*. This representation (with scalar reward values associated with each state) was used in early work to model temporal difference learning in tasks such as the example considered here (Schultz et al., 1997). An immediate problem with this model is that it does not represent the ITI, an interval that has been shown repeatedly to affect conditioning (Gibbon and Balsam, 1981; Holland, 2000; Kalmbach et al., 2019; Lattal, 1999). The ITI is almost always a random variable with a specified probability distribution. Since Markov chains assume that *all the states* must be specified, there is no obvious way to break up the ITI into a fixed set of equal duration states obeying the Markov property. This problem is usually avoided by only modeling the “trial period”, i.e. the delay between cue and reward. However, this is evidently an incomplete representation of the task, the stated goal of a state space. Nevertheless, this model has proven to be quite successful at explaining numerous aspects of conditioning and thus, has been referred to as a “useful fiction” (Ludvig et al., 2012; Sutton and Barto, 1998).

**Fig 2.**
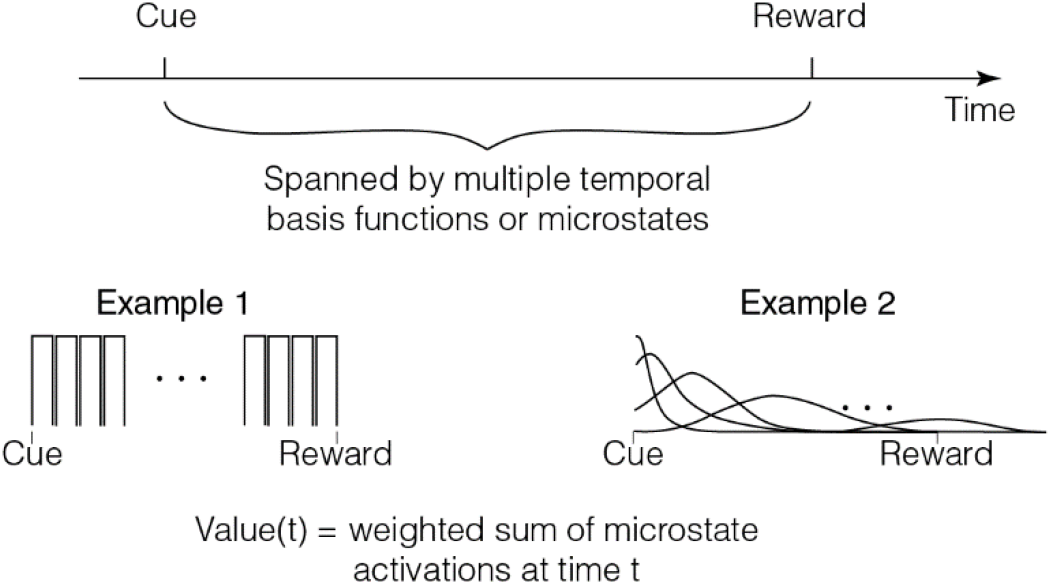
Common models for dealing with delays between cue and reward assume that such delays are spanned by multiple microstates. Two examples are shown here (see text). As can be seen, these formulations assume that the delay periods themselves are represented by many states to which an RL algorithm can attach value.

An extension of this model is to treat the states in the delay not as fixed duration states, but as a set of basis functions (also known as microstates or microstimuli) (Ludvig et al., 2008, 2012) (example 2 in **Fig 2**). A convenient idea is that the delay after a cue is spanned by a consecutive set of Gaussian states (Ludvig et al., 2008, 2012). In this view, each subsequent state has progressively smaller amplitude and larger width (to approximate scalar timing). This model of state space has benefits over the complete serial compound, as it allows efficient generalization and flexibility due to the non-zero value of many microstates at any given moment (Gershman et al., 2014; Ludvig et al., 2008, 2012; Petter et al., 2018). There is also some evidence for microstate-like activity patterns in brain regions such as the striatum (Mello et al., 2015), hippocampus (MacDonald et al., 2011; Pastalkova et al., 2008; Salz et al., 2016), and the prefrontal cortex (Tiganj et al., 2017). Remarkably, these neural representations can flexibly scale when the delays are altered (MacDonald et al., 2011; Mello et al., 2015). Thus, this microstate model is more consistent with neural data and is functionally advantageous over the complete serial compound. There are many variants of this general idea of a series of microstates (i.e. sequential set of delay states) following a cue (Brandon et al., 2003; Desmond and Moore, 1988; Grossberg and Schmajuk, 1989; Machado, 1997; Mondragón et al., 2014; Vogel et al., 2003; Wagner, 1981). We will not review these in detail here.

## Implicit assumptions

The fundamental premise of the above models is that the delay between different environmental stimuli is a sequence of states in an animal’s state space. By breaking the flow of time into such sequences of states, these models make some implicit assumptions. These are often not immediately obvious. We will list some here.

1. <u>Every cue has its own associated set of microstates:</u> the idea of microstates works only if separate cues have separate sets of microstates. Thus, if the animal is learning that cue1 predicts reward1 after delay1 and cue2 predicts reward2 after delay2, the set of microstates during delay1 must be different from the set of microstates during delay2. If not, value learning will be mixed up between the two cues and cannot appropriately assign credit.
2. <u>The microstates are specified *before* value learning:</u> this is may be the most important assumption. The entire idea of RL (with value updates to satisfy the Bellman equation) works only if the state space is specified. Thus, before value learning can occur, the set of sequential microstates following a cue must already exist. We will discuss the problems with this assumption in more detail in the next section.
3. <u>Number and form of microstates are free parameters:</u> another major assumption is that the number and form (e.g. are the basis functions Gaussian?) of microstates are treatable as free parameters for model-fitting. While the lack of principles for the formation of microstates is an obvious problem, this assumption is especially problematic as conditioning in the laboratory can occur over delays of milliseconds or even twenty four hours (Etscorn and Stephens, 1973; Hinderliter et al., 2012; Kehoe and Macrae, 2002). It is unclear what, if any, principles govern the formation of microstates in the brains of real animals over spans of five orders of magnitude.
4. <u>The microstates during the cue to reward delay are fundamentally different from the microstates during the ITI:</u> In the microstate framework, different delay periods that contain no external stimuli must be treated differently. Thus, the set of microstates during the cue to reward delay must be different from the set of microstates from the reward to cue delay. An explicit treatment of this formulation is found in the SOP model (Wagner, 1981).
5. <u>Learning occurs in trials:</u> another implicit assumption is that value learning occurs progressively by accumulation across trials. It is this trial duration that is assumed to be split into microstates. However, experiments such as the truly random control (Rescorla, 1967, 1968) throw the validity of this assumption into question. In this experiment, cues and rewards are both delivered by independent Poisson processes and hence, have no statistical relation to each other. Worse, because the events are Poisson processes, they are equally likely to occur at any moment in time. In this case, it is unclear what, if anything, can be treated as a “trial” in the animal’s brain.
6. <u>Microstates of a cue must be reproducible across repeated presentations:</u> for learning to occur, if cue1 evokes a set of microstates on one presentation, the same set of microstates must be evoked on the next presentation, to ascribe value to the “correct” microstate.

In the next section, we take a deeper dive into these assumptions and show that the apparent simplicity of the microstate model belies a gargantuan complexity of representation imputed in animal brains. We are by no means the first to discuss some of the problematic implications of these assumptions (Gallistel et al., 2014, 2019; Hallam et al., 1992; Hammond and Paynter Jr, 1983; Luzardo et al., 2017). Nevertheless, the following section focuses on a particularly problematic aspect of these assumptions that has not received as much discussion in the literature.

## How bad is the problem really?

The problem is brought into sharp relief when considering initial learning. Remember that the whole point of the formulation of a state space is to explain reward prediction learning. Thus, we will now critically examine the implications of these assumptions for initial learning.

Imagine an animal that is first experiencing a cue that will be followed later by a reward. On this first experience, the animal knows nothing of the significance of this cue (other than its general “salience” or intensity). Indeed, cues are galore in the environments of animals. Nearly every sensory feature of the world could in principle be a cue predictive of a future reward. For instance, may be a sound is predictive of a future reward. If an animal indeed learns to predict this reward, the above RL algorithms would require the assumption that the sound evokes microstates until the reward *before first learning the relationship of the sound to reward*. This is the whole point of RL: it is a model of learning after all.

What does this imply? This implies that any cue that could *in principle* be predictive of reward must evoke microstates during every presentation. Every sensory stimulus could in principle be such a cue. Hence, for the microstate model to work, animal brains must produce microstates for every sensory stimulus in the experience of the animal. Worse, if cue1 was experienced on two separate days, the set of microstates that were evoked by cue1 should be the same. Thus, the brain must store in memory all the microstates for the nearly infinite number of sensory stimuli and they must all be discriminable and reliably reproducible on repeated presentations of the stimuli.

The problem is actually much worse. This is because the animal does not know at what delay a reward will follow a cue on the first experience of the cue. Indeed, these delays can span five orders of magnitude (Etscorn and Stephens, 1973; Hinderliter et al., 2012; Kehoe and Macrae, 2002). As mentioned above, the data that are often taken as evidence of the existence of neural microstates show that these time representations remap when the delay changes (MacDonald et al., 2011; Mello et al., 2015). How then does the brain know what exact microstates to trigger on the first presentation of the cue, much before the delay to reward is known or learned? Worse still, the brain also must trigger microstates during the delay from the reward to the next cue, for every reward and cue, to learn the distribution of intertrial intervals. How does the brain produce distinct microstates during the ITI and delay to reward on the *first presentation of the cue and reward*? How does the brain know that two delay periods during which no external sensory stimuli exist are fundamentally different *before learning* that there is a relationship between cue and reward? How also does the brain know that delays between different cue-reward-cue pairs are different? It is hopefully clear from this examination that the assumption of microstates, while seemingly simple, introduces an untenable solution for an animal brain. Solving these issues is crucial as these issues riddle application of RL to animal learning in even one of the simplest use cases considered here.

One approach that has proven quite successful at explaining numerous timing phenomena related to conditioning is the combination of a Rescorla-Wagner rule applied to a drift diffusion model of timing (Luzardo et al., 2017). In this model, cues are postulated to initiate an accumulating timer with a fixed threshold and an adaptable slope. A learning rule adapts the slope based on the knowledge of when the reward happens, thereby adapting the slope of the accumulator to eventually time the cue-reward delay appropriately. This model explains an impressive array of phenomena. It also has a major advantage over the microstate models as it does not postulate an arbitrary number of microstates that span time delays. Nevertheless, it too suffers from similar issues as above when applied to initial learning. For it to work for initial learning, there must be a timer for every cue that could in principle be predictive of reward. As we laid out above, there are almost an infinite number of such cues. Further, when a timer is initiated at cue onset on the first time that the cue was experienced, how does the timer know that it is timing a specific upcoming reward? What if this cue was only predictive of another cue and not a reward? How does the timer get feedback about exactly which interval it is supposed to time? These issues are solvable if the animal knows that it is timing the interval between a specific cue state and a reward state, or in other words, *after* learning that a specific cue and a specific reward may be related.

## Proposed solution: continuous time Markov renewal process state space

One possible solution to the problem is to move completely away from RL and propose that other quantities control learning in tasks such as the one considered here. A set of models that propose that animals learn contingency (defined as normalized gain in available information) between the *timing* of reward predictors and rewards belongs to this class (Balsam et al., 2010; Gallistel et al., 2014, 2019; Ward et al., 2012). These models are successful at explaining numerous aspects of the learning of conditioned responses in relation to the various time intervals. Further, they can work for initial learning in a timescale invariant fashion. Here, we propose an alternate model for the state space representation in the tradition of RL. To this end, we use a state space based on a Markov renewal process. While this proposed model needs to be empirically tested, we show that it at least does not contain the same problems as contemporary microstate models, can naturally incorporate the continuous passage of time, and be applicable to initial learning. Our formalism is an extension of previous semi-Markov RL models (Bradtke and Duff, 1994; Daw et al., 2006).

In our model, the set of states in the cue-reward task are treated as *S = {cue, reward}*. No delay periods are treated as their own states. For simplicity, we will assume that these states can be described by the events denoting their onsets. This is not a requirement of the formalism and we do so purely for simplification of this treatment. Given this assumption, we can represent the experience of an animal using a timeline (**Fig 3A**). Here, the delays between state onsets (i.e. events) can take any value and has a specified distribution. For instance, if the reward follows the cue at a fixed delay, the interevent distribution is a Dirac delta function at this delay. In general, the transition properties of such a sequence of states is fully described by the probability “kernel” shown in **Fig 3**. A dynamic estimate of value for continuous time can be obtained for such a representation, based on the intuition that animals maximize quantities such as reward rate (Blanchard et al., 2013; Gallistel and Gibbon, 2000; Hamid et al., 2015; Namboodiri et al., 2014a; Stephens and Krebs, 1986). We mathematically derive this estimate in Appendix 1 based on the transition kernels shown in **Fig 3**. We further describe approximations to estimate reward rate that are consistent with several well-established observations in animal behavior (see *Implications* below).

**Fig 3.**
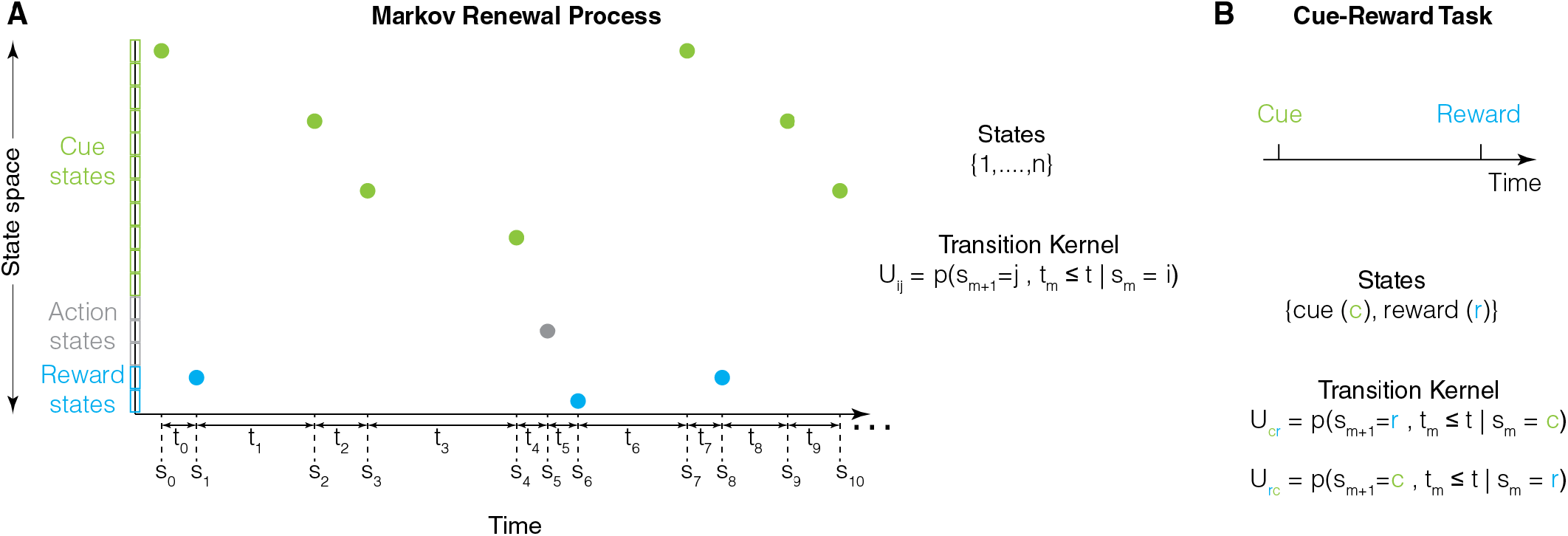
**A.** A natural means to represent the timeline of experience of animals is by using a Markov renewal process. Here, observable events (and not delays between them) are the states. In the timeline, there is a sequence of states (denoted by s_0_,…,s_10_,…) and transition times between them (t_0_,…,t_9_,…). The state at each transition can be one among the set of states denoted by {1,…,n}. Given this, the evolution of the states can be defined by a transition kernel *U*_*ij*_, which measures the joint probability of state identity and cumulative transition time for the next transition. **B**. The above formalism naturally accounts for the cue-reward delay (in *U*_*cr*_) and the intertrial interval (in *U*_*rc*_) (Appendix 1).

In this continuous time model, only observed events (i.e. cue and reward) are treated as their own states on the first exposure of these stimuli. Thus, there is no requirement of arbitrary microstates to be present during delay periods prior to initial learning. Any organism that stores its experience in a timeline (e.g. **Fig 3**) can perform the computations necessary to estimate the continuous time rate contingency and value described here (see Appendix 1 for details). Recent evidence suggests that such timelines exist in animals (Bright et al., 2020; Panoz-Brown et al., 2018; Tiganj et al., 2018; Tsao et al., 2018; Zhou and Crystal, 2009). Thus, we believe that this algorithm defines a state space that is suitable for real animals and is a natural extension of reinforcement learning to continuous time.

## Implications

The fundamental premise of our model is that animals do not discretize or break up the flow of time. Our proposal is one possible solution that can handle the continuous passage of time in learning the state space and contingency. There may well be others and our hope here is to move the field forward by explicitly listing out the problematic implications of well-known neuroscience-related RL models in relation to their handling of time delays. Indeed, our model is philosophically a hybrid between the semi-Markov formulation of state space (Bradtke and Duff, 1994; Daw et al., 2006), and the contingency of timing model based on information theory (Balsam and Gallistel, 2009; Balsam et al., 2010; Gallistel et al., 2014). The latter model proposes that animals compute the information gained from a cue on the timing of upcoming rewards. This model has not as yet been extended to sequences of states or stimuli that predict reward, and does not provide a clear explanation for the error prediction signals observed in midbrain dopamine neurons (Cohen et al., 2012; Kim et al., 2020; Mohebi et al., 2019; Schultz, 2016; Schultz et al., 1997). On the other hand, as our model is built on learning state transitions, it retains the natural scalability of RL formulations to extended sequences of states predictive of reward. It also retains the possibility of error prediction learning on these states. These extensions are beyond the scope of this paper. Our model can also in principle be extended to continuous state spaces using previous approaches (Doya, 2000).

Our proposed solution is consistent with a few well-established facts in animal behavior that violate predictions of commonly used RL algorithms in neuroscience. First, an approximate solution to the Markov renewal process rate estimation (exact equation in A1.7) results in hyperbolic discounting (as shown in A1.23). This contrasts with the exponential discounting commonly assumed in reinforcement learning algorithms (Sutton and Barto, 1998). Observations from real animals (including humans) convincingly demonstrate hyperbolic discounting, as have been reviewed previously (Frederick et al., 2002; George Ainslie, 2001; Namboodiri et al., 2014a). The approximation in A1.23 is also consistent with numerous related aspects of temporal decision-making (Namboodiri et al., 2014a). Second, our framework predicts that animals calculate the mean transition time from the current state to all possible future states (equation (A1.21)). Consistent with this, prior research has observed “temporal averaging” of the expected delay to reward from multiple cues (De Corte and Matell, 2016; Matell and Henning, 2013; Matell and Kurti, 2014). Third, our framework implies that initial learning does not proceed by the gradual accumulation of reward prediction or weights (e.g. Rescorla-Wagner or temporal difference learning). Instead, we propose that learning proceeds by the computation of a reward rate contingency (Appendix 1). Such a computation implies a delay until the contingency is deemed to cross a statistical threshold. Thus, our model implies a sigmoidal learning curve instead of an exponential learning curve, an implication well-supported by data from individual subjects (Gallistel et al., 2004; Morris and Bouton, 2006; Pamir et al., 2011, 2014; Papachristos and Gallistel, 2006; Takemoto et al., 2015; Ward et al., 2012). Lastly, as expected from equation (A1.24), the number of reinforcements to such acquisition depends positively on the intertrial interval and negatively on the cue-reward delay (Gibbon and Balsam, 1981; Holland, 2000; Kalmbach et al., 2019; Lattal, 1999). This prediction is qualitatively similar to that made by the contingency of timing models, and can also be accounted for by other continuous time models (Luzardo et al., 2017; Shankar and Howard, 2012).

In light of some qualitative similarities between our model and the contingency of timing model, we will now consider some of their differences. One distinction relates to distractor states. Our model is predicated on the learning of transitions between adjacent states (Equation (A1.1)). If during learning a cue-reward association, animals also experience salient distractor cues in the delay periods that get treated as states, we predict that the learning will at least be temporarily disrupted. This is because the next state from the cue state will often be the distractor state. To learn the relationship between cue and reward, the animals will need to learn to omit the distractor state from the state space of the task. In contrast to this prediction, the contingency of timing model predicts that distractor states will have no effect. This is because it postulates that animals estimate the delay from every cue state to the next reward state *regardless of any intervening states*. Existing evidence supports the prediction that distractor states impede trace conditioning (Carter et al., 2003; Clark et al., 2002; Han et al., 2003; Manns et al., 2000).

These data raise the question of what events get treated as states during a task. While saliency is likely to play a major role (e.g. barely audible noise is less likely to be treated as a state than a loud noise), it may well be that animals experience *subjective* states during delays (i.e. sates that were not part of the experimental design). The likelihood that such subjective distractors affect conditioning will increase with increasing delay periods. This is because the longer the delays, the higher the chance that animals experience an internal subjective state independent of the task design. Hence, another qualitative difference between our model and the contingency of time model is that with increasing delays, the scalar relationship seen in Equation (A1.24) may no longer hold due to the presence of subjective distractor states that impede contingency learning. This may explain why in some appetitive conditioning protocols, no learning is seen for minute long trace intervals, even when intertrial intervals are very long (Thrailkill et al., 2020). An alternative possibility is that in these cases, the look-ahead time *T* is not significantly longer than the cue-reward delay, thereby reducing reward rate contingency or value (Equation (A1.24)).

Another implication of our model is that the brain must be capable of storing recent memory in a timeline of events, based on which the calculation of contingency described above may be performed. This implication is shared by the contingency of timing models. Behavioral evidence suggests that animals are capable of storing past experiences in a timeline (Panoz-Brown et al., 2016, 2018; Zhou and Crystal, 2009). A Laplace-transform based computational model provides one potential neural network solution for storing recent history in a timeline (Shankar and Howard, 2012). Recent evidence shows that the entorhinal cortex or prefrontal cortex can store temporal information in learned memories (Bright et al., 2020; Tiganj et al., 2018; Tsao et al., 2018), although the extent to which these systems act as an accurate organizer of timestamps of recent memories (similar to **Fig 3**) remains to be tested. Considering the memory requirements of storing such timelines, another intriguing possibility is that the brain has an explicit memory medium inside neurons (Akhlaghpour, 2020; Langille and Gallistel, 2020). Lastly, another common assumption of our model and the contingency of time model is that a contingency calculation can be learned in principle for all cues predictive of reward. Considering the computational requirement of such updating, it is unknown how the brain might solve this problem. An intriguing solution for this problem was discussed recently using a representation of retrospective transition probabilities *p(s*_*m*_=*cue*|*s*_*m+1*_=*reward)* (Namboodiri et al., 2019). The corresponding prospective probabilities may be learned by Bayes’ inversion of these retrospective probabilities. Considering the ethological sparsity of rewards, the updates of the retrospective probability, triggered only on reward receipt, are much sparser. Experimental data show that some neuronal subpopulations in the orbitofrontal cortex show activity patterns consistent with such retrospective representations and affect behavioral learning and memory (Namboodiri et al., 2019).

Finally, for simplicity, we have illustrated our main point using the simplest form of RL—one in which the selection of actions to maximize future rewards is not considered. The consideration of time delays becomes even more important for action selection. For instance, it is common in RL to describe reward prediction by either a dual conditional probability *p(reward|state,action)* or a dual conditional reward function *R(state, action)*. Real animals often perform reward-related actions after a delay from the corresponding reward-related cues. Indeed, rewards are often predicted by (cue, action) pairs only when there is a specific temporal relation between these events (e.g. Miyazaki et al., 2020; Namboodiri et al., 2015; Narayanan and Laubach, 2009). Defining microstates to span these delays worsens the combinatorial explosion of the state space, as these microstates need to then depend on both external cues and internal actions. Thus, the issues discussed here become even more vexing in this setting.

## Conclusions

Here, we explicitly list the assumptions made by the well-known RL models that account for the passage of time. We show that the apparent superficial simplicity of these models belies the extraordinary complexity required to execute them. These problems are often not recognized, as researchers define a convenient state space for each experiment using free parameters. These assumptions are especially problematic when applied to initial learning, a stated goal of reinforcement learning. Our hope is that the field of neuroscience-related RL explicitly contends with the issues we have raised and moves our knowledge forward by defining the state space of real animals.

## Acknowledgments

I thank J. Berke, C.R. Gallistel, S. Mihalas, and members of the Namboodiri lab for helpful comments that instructed this formalism and the manuscript. This work was funded by grants from the National Institute of Mental Health (R00MH118422), Brain and Behavior Research Foundation (NARSAD Young Investigator Award), and the Scott Alan Myers Endowed Professorship to V.M.K.N.

## APPENDIX

## Appendix 1: Mathematical treatment of rate estimation in a Markov renewal process

The transition probability kernel for a Markov renewal process is expressed as

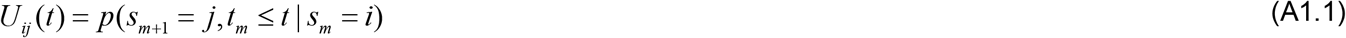

Where *s*_*m*_ is the m^th^ state in a sequence and *t*_*m*_ is the transition time for the m^th^ transition. Thus, this kernel is the joint probability of transitioning from state *i* to *j* within a delay of *t*. In general, the delay distribution can be arbitrary.

Given this formalism, the key step now is to postulate the goal of learning. In standard reinforcement learning, the goal is taken to be to estimate the value function (Sutton and Barto, 1998). The value function could in principle be defined as the sum of all rewards obtainable in the future, given the current state. However, since this sum does not converge, it is common to assume an exponential discounting of future rewards based on the number of timesteps to those rewards (Schultz et al., 1997; Sutton and Barto, 1998). This assumption is primarily made for mathematical simplicity, as it allows for a recursive definition of value between successive timesteps. Despite this simplicity, behavioral evidence from animals convincingly demonstrate that this assumption is false (Frederick et al., 2002; George Ainslie, 2001; Namboodiri et al., 2014a).

Instead of arbitrarily assuming exponential discounting, we define the general goal of learning for animals as the estimation of the rate at which a state *j* will be visited in the future (say within a look-ahead time of *T’*) if the current state is *i*. The use of event rate is similar to many prior proposals (Gallistel and Gibbon, 2000; Namboodiri et al., 2014a; Stephens and Krebs, 1986). For reward prediction, state *j* can be thought of as the reward state and state *i* as a reward predictor state (e.g. cue state). We will denote this expected rate by *λ*_*ij*_*(T’).* In general, if this rate is different from the rate of visits to state *j* from a random moment in time (denoted by *λ*_*−j*_*(T’)*, where the - in the subscript denotes a random moment in time), then state *i* is predictive of state *j* in the future. This is because *λ*_*ij*_*(T’)* > *λ*_*−j*_*(T’)* implies that starting in state *i* will result in more future visits of state *j* than starting at a random moment in time. Hence, the fundamental premise here is that animals learn both *λ*_*ij*_*(T’)* and *λ*_*−j*_*(T’)*, and that their difference measures the “rate contingency” of state *j* on state *i*. Another intuitive way to think of these rates is as the conditional rate of visiting state *j* conditioned on starting in state *i* minus the marginal rate of visiting state *j*. When the conditional equals the marginal, state *i* does not predict the future occurrence of state *j* beyond chance.

*λ*_*ij*_*(T’)* can be calculated by first calculating the expected number of times state *j* will be visited within a future look-ahead time of *T’* by starting in state *i*. We will denote this number by *M*_*ij*_*(T’)*. This estimated number is the continuous time equivalent of the undiscounted successor representation in discrete-time Markov chains (Gershman, 2018; Momennejad et al., 2017; Russek et al., 2017), as explained below.

In discrete time, the expected number of times that a Markov chain will visit state *j* in the future within *t* time steps is

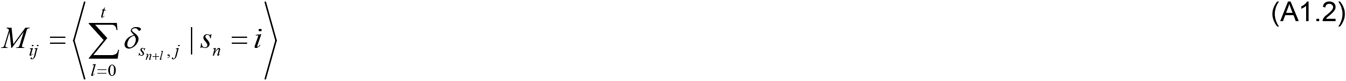

Where *δ*_*ij*_ is the Kronecker-delta function which is one when its arguments are equal, and zero otherwise. Thus, the *δ* in the sum is 1 when a future state equals j and 0 otherwise.

Since this sum does not converge as *t* becomes very large, it is customary to assume a discounting for future visits that are further ahead in time. If an exponential discounting is assumed at a rate of *γ* for every time step, then a modified version of the above sum for the discounted occupancy of state *j* in the future starting from state *i* in the present is

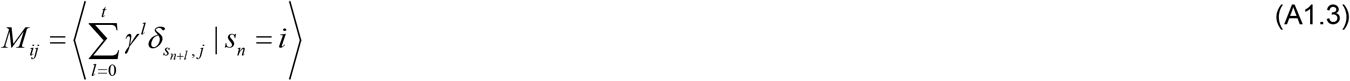

This is the definition of the successor representation for a discrete time Markov chain (i.e. without actions) (Gershman, 2018; Momennejad et al., 2017; Russek et al., 2017).

The continuous time equivalent of the undiscounted sum (similar to (A1.2)) in a Markov renewal process is the expected number of times state *j* will be visited in within a future look-ahead time of *T’* by starting in state *i*. Thus, the rate of occurrence of state *j* is

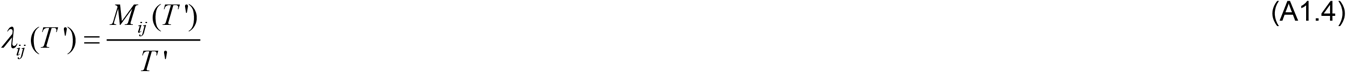

Once learned, the above estimate of *λ*_*ij*_*(T’)* can be used to calculate the expected reward rate by starting in state *i* in the look-ahead *T’* by the following calculation

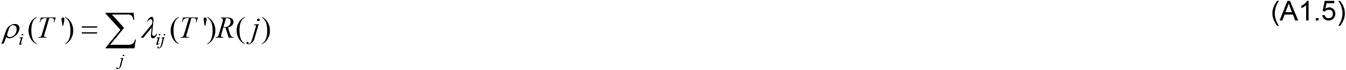

Where *R(j)* is the reward associated with state *j*.

The difference between the above estimate of reward rate and the average reward rate of starting at a random moment in time (denoted by *ρ-)* provides an estimate of the value of starting in state *i*.

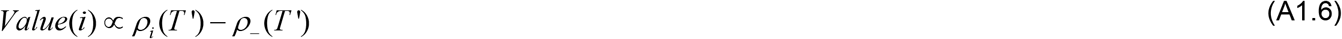

Given these relationships, the animal needs to learn *M*_*ij*_*(T’)* to calculate the value of being in state *i.* As mentioned above, *M*_*ij*_*(T’)* is the expected number of times that state *j* will be visited within a look-ahead time of *T’* by starting in state *i*. This can be calculated by first counting whether state *i* (the starting state) is itself state *j* (in which case the count should increase by 1), and then by estimating the expected total number of times state *j* will be visited from any state *k* to which state *i* transitions, in the remaining time after the transition from *i* to *k*. Mathematically, this is written by the following integral equation.

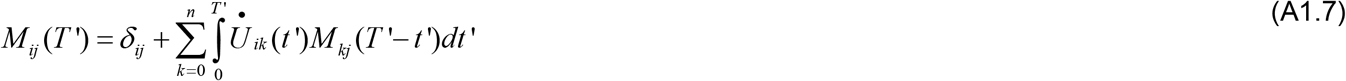

Where 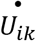 is the time derivative of the transition probability kernel (representing the instantaneous probability density of transition between *t’* and *t’+dt’*).

This integral equation provides the exact solution for *M*_*ij*_*(T’)*, which can be substituted in equation (A1.4) above to calculate the value of *λ*_*ij*_*(T’)*. We can extend equation (A1.7) to obtain a dynamic estimate for *M*_*ij*_*(T’)* after a time *t* has elapsed without a transition following entry into state *i*. If we represent this by *M*_*ij*_*(T’, t)*, we can calculate its value by

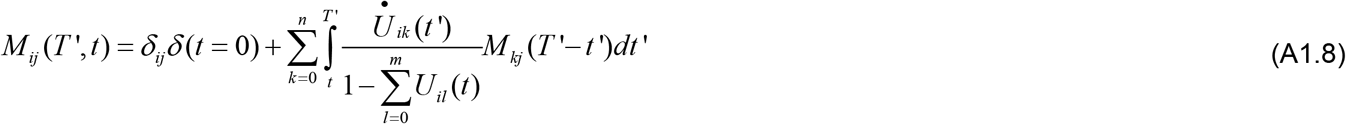

Here, *δ(t=0)* is the Dirac delta function, which is 1 when *t=0* and zero otherwise (i.e. increase expected count of *j* by 1 at onset of state *i* if state *i* is the same as state *j*.

The ratio 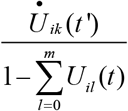 represents the instantaneous probability of the transition to state *k* from *i* between *t’* and *t’+dt’*, given that no transition occurred between *t’=0* and *t’=t* following entry into state *i*. This conditional probability is the ratio of the joint probability divided by the probability of the conditioned event. The joint probability is the same as the instantaneous probability of transitioning from state *i* to state *k* between *t’* and *t’+dt’*, as *t’>t*. The probability of the conditioned event is the probability that no transition occurred between *t’=0* and *t’=t* following entry into state *i*, and this is the denominator in the ratio.

Based on these equations, learning the estimated rate *λ*_*ij*_*(T’)* or *λ*_*ij*_*(T’, t)* can be achieved through several potential means. We do not discuss this here since our primary objective is to convey that there is a natural means to define a value function for continuous time, based on a reward rate contingency.

Solving equations (A1.7) or (A1.8) is challenging. An exact method could be based on explicit counting on a timeline such as that shown in **Fig 3**. Here, we will propose three approximate methods to solve equation (A1.7) to derive interpretable predictions.

## Approximation 1

Before learning *U*_*ik*_, one approximation is to simply assume that the second term above is the average rate of occurrence of state *j* from a random moment in time. In this case,

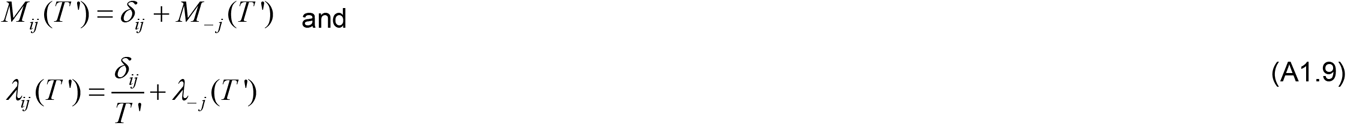

Therefore, the rate contingency is simply 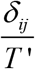.

## Approximation 2

Once *U*_*ik*_*(t)* has been learned, *M*_*kj*_*(T’-t’)* can be expressed as

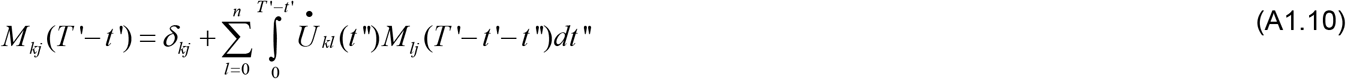

The second term on the right-hand side (RHS) of this equation can be approximated by the expected value of *M_-j_(T’-t’)*. Essentially, in this approximation, we assume that after the first transition from *i* occurs to *k*, the remainder of the transitions to *j* can be approximated by counting whether *k* is *j* and then estimating the baseline expected number of transitions to *j* in the time period *T’-t’*. Thus, equation (A1.10) can be approximated to

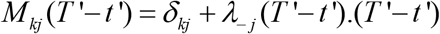

Further, assuming *T’>>t’*, the baseline rate *λ*_*−j*_*(T’-t’)* can be assumed to be independent of the look-ahead period and rewritten as *λ*_*−j*_.

Therefore, we get the following approximation.

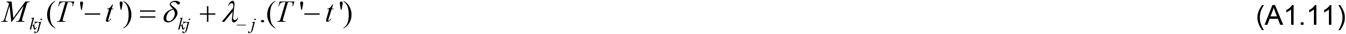

Substituting (A1.11) in (A1.7), we get

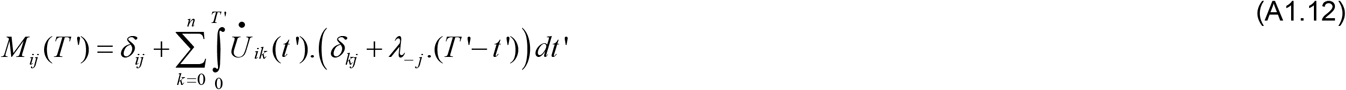

The second term of the RHS of this equation can be split into two terms. We will now derive the simplified values of these terms.

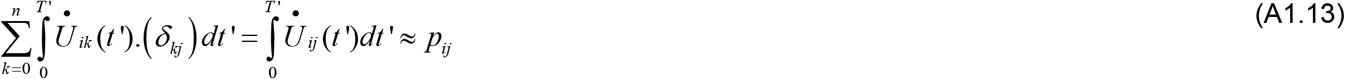

Here, we assumed that *T’* is large enough that the integral can be approximated by its value integrated to the upper limit of ∞. *p*_*ij*_ is the probability of transition from state *i* through state *j* over an infinite time, i.e. the transition probability of the underlying Markov chain.

The next term in the RHS of equation (A1.12) is

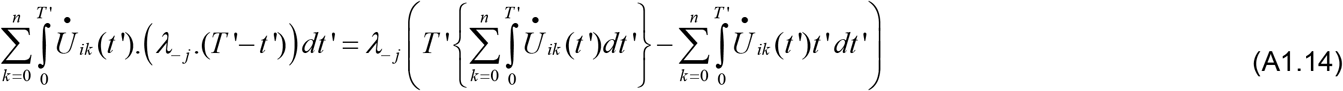

Again, if *T’* is large enough, the first term in the curly parenthesis in the RHS of (A1.14) can be approximated as

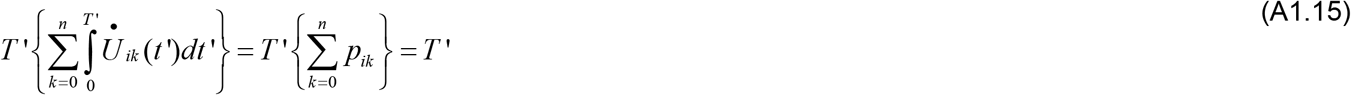

The second term in the RHS of (A1.14) can similarly be approximated as

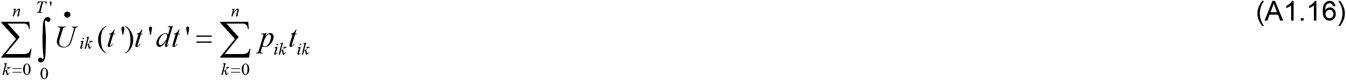

Where *t*_*ik*_ is the mean transition time between state *i* and state *k*.

Substituting (A1.15) and (A1.16) in (A1.14), we get

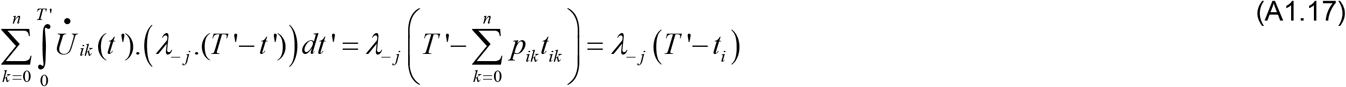

Where *t*_*i*_ is the mean transition time from state *i*.

Substituting (A1.17) and (A1.13) in (A1.12), we get

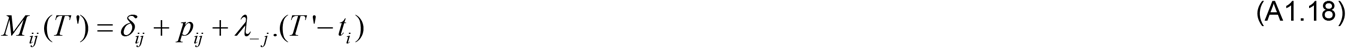

Therefore, the rate *λ*_*ij*_*(T’)* is

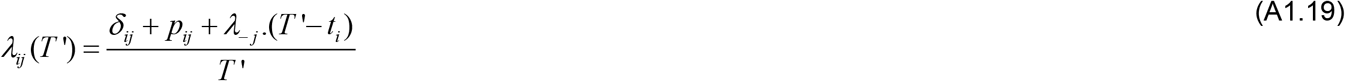

If we now assume that the look-ahead *T’=T+t*_*i*_, where *T* is the amount of look-ahead beyond the average transition time from the current state *i*, then (A1.19) can be rewritten as

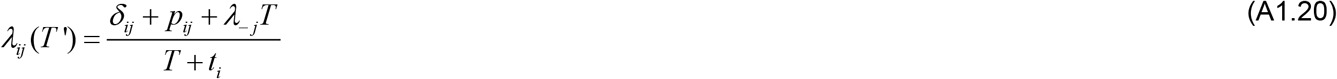

This is the approximate value for *λ*_*ij*_*(T’)*= *λ*_*ij*_*(T+t*_*i*_). Essentially, we assume that the look-ahead time period is *added* onto the expected transition time from the current state *i*.

Therefore, the rate contingency is

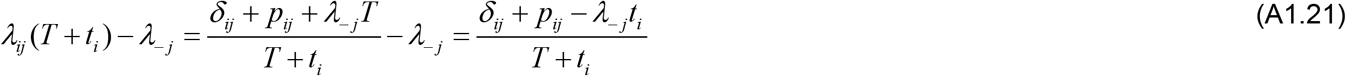

The appearance of the average transition time from the current state (*t_i_*) in the above equation suggests that animals may estimate this interval. Consistent with this, prior research has observed “temporal averaging” of the expected delay to reward from multiple cues (De Corte and Matell, 2016; Matell and Henning, 2013; Matell and Kurti, 2014).

For a task in which a cue (state *i*) is followed by a reward (state *j*) at a fixed delay *t*_*r*_ with a probability *p*_*r*_ and the intertrial interval denoted by *t*_*ITI*_, *p*_*ij*_=*p* and *t*_*i*_=*pt*_*r*_+*(1-p)(t*_*r*_+*t*_*ITI*_). Substituting these, we get

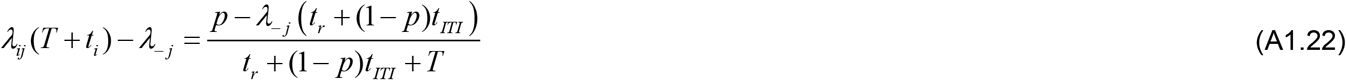

For a reward *r* available with 100% probability, the subjective value—defined as the reward delivered with zero delay that is treated subjectively equivalent to the delayed reward—is therefore

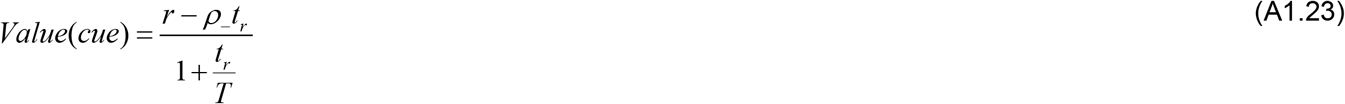

This relationship shows that delayed rewards are hyperbolically discounted, with the rate of discounting determined by the opportunity cost of losing rewards during the wait to reward (in the numerator), and by the look-ahead time period *T* (in the denominator). This is a special case that has been described in detail previously and fits numerous experimental observations related to delay discounting (Namboodiri et al., 2014b, 2014c, 2014a, 2016a, 2016b). In keeping with this prior set of publications, we will refer to this approximation of the Markov renewal process as the TIMERR approximation.

If the marginal reward rate *ρ-* has been appropriately learned in a stationary environment, its value should be the reward obtained after the cue divided by the total trial duration (sum of delay from cue to reward, and the intertrial interval). In this stationary case, the subjective value of a cue paired with delayed reward is

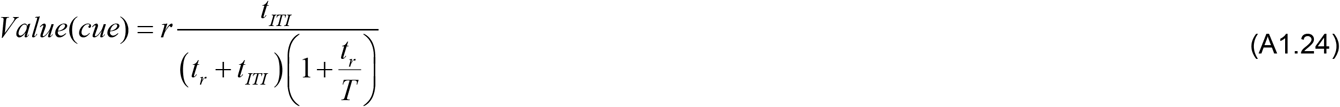

When *T>>t*_*r*_, i.e. the look-ahead time is much greater than the cue-reward delay (the basis for the above approximations), the above relationship is scalar with respect to the intertrial interval and the cue-reward delay. In other words, scaling up both these intervals by the same amount will have no effect on the value of the cue.

Overall, the above results show that a continuous time model of learning and decision-making with a dynamic estimate of value can be naturally defined using a Markov renewal process (equations A1.1, A1.4–1.8). Simplifying approximations to obtain interpretable results show both hyperbolic discounting (equation (A1.23)) and a scalar relationship of the intertrial interval and cue-reward delay (equation (A1.24)), as have been experimentally observed (Blanchard et al., 2013; George Ainslie, 2001; Gibbon and Balsam, 1981; Holland, 2000; Kalmbach et al., 2019; Lattal, 1999).

## Approximation 3

Another possibility is to potentially approximate *M*_*kj*_*(T’-t’)* as

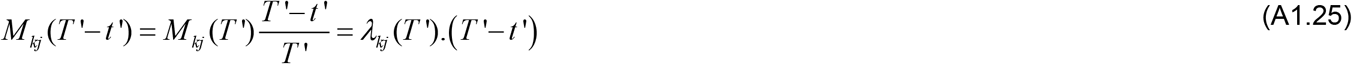

This approximation essentially assumes that *λ*_*kj*_ is independent of the remaining look-ahead time period. In other words, this approximation is the same as assuming a constant rate of events.

Substituting (A1.25) in (A1.7), we get

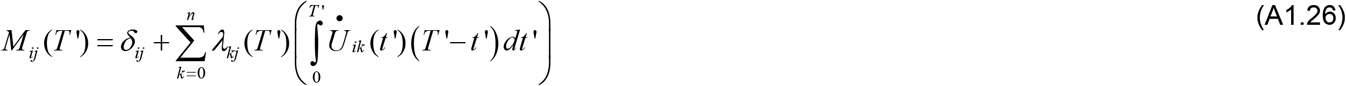

If *T’* is large enough, similar to the approximations used in (A1.15) and (A1.16), the term in the parenthesis can be approximated as

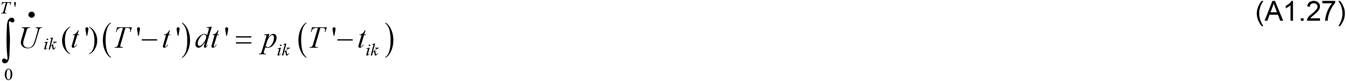

Substituting (A1.27) in (A1.26), we get

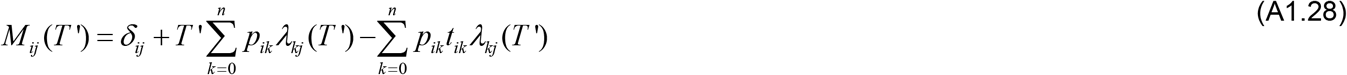

Thus, *λ*_*ij*_*(T’)* is

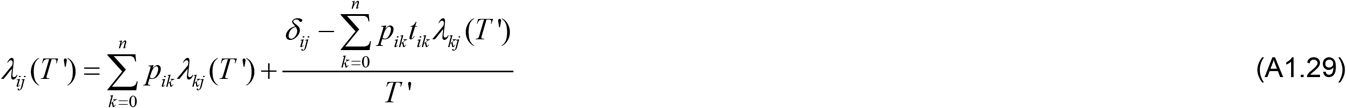

If the matrices representing *λ_ij_*, *p_ij_* and *t_ij_* are ***λ, P*** and ***t*** respectively, the above equation can be written as

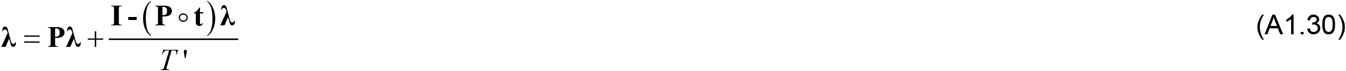

Where ∘ represents the Hadamard or Schur or element-wise product. Solving the above matrix equation for ***λ***, we get

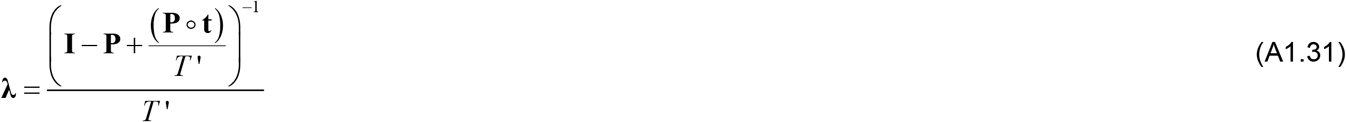

For the simple cue-reward task, we can write

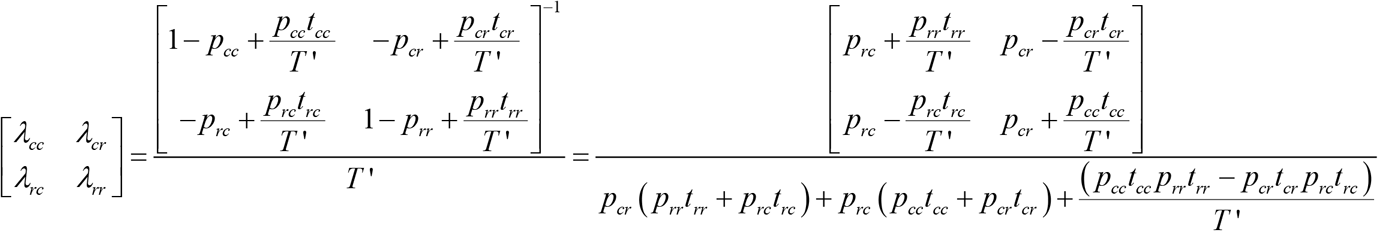

Setting *p*_*cr*_=*p*_*r*_, *t*_*cr*_=*t*_*r*_, *p*_*rc*_=1, *t*_*rc*_=*t*_*ITI*_, *p*_*cc*_=*1-p*_*r*_, *t*_*cc*_=*t_r_*+*t*_*ITI*_ and *p*_*rr*_=*0*, we get

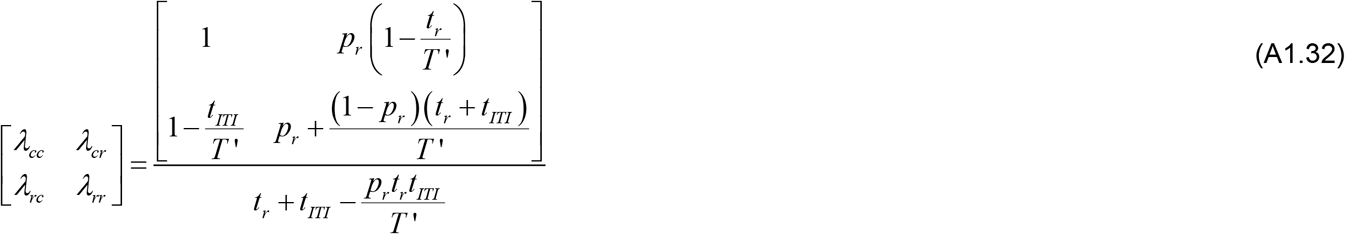

Here too, for the large value of *T’* assumed, we obtain the estimated rate of rewards following cue of *p*_*r*_/*(t*_*r*_ + *t*_*ITI*_). Thus, for an environment that is assumed to be completely stable in the future, starting in the cue state is the same as starting at any random moment in time, as the expected rate of rewards over an infinite time horizon is the same. Since animals are unlikely to assume such stationarity, approximation 2, i.e. the assumption that the rate of rewards beyond the next transition is equal to the base rate, is more appropriate. This model is also more consistent with animal behavior, since it is consistent with the hyperbolic discounting and opportunity costs observed in animal behavior (Namboodiri et al., 2014b, 2014c, 2014a, 2016a, 2016b, 2019).

Overall, these approximations provide potential means to calculate rate of rewards using a continuous time formalism. How animals learn these quantities is beyond the scope of this paper. Nevertheless, we will mention two possible alternatives. A model-free learning rule for estimating value may be defined based on equation (A1.6) by directly estimating rates of rewards in a look-ahead period from cue onsets. In a model-based conception, the transition kernel *U_ij_* shown in equation (A1.1) needs to be learned. Considering the computational cost of the model-based method, it may be possible that initial learning is model-free, and when the value of a cue is deemed high, the brain calculates an explicit model-based value. Another possibility is that initial learning is driven not by estimates of prospective transition probabilities, but by estimating retrospective transition probabilities—a considerably sparser learning rule (Namboodiri et al., 2019).

## References

Akhlaghpour, H. (2020). An RNA-Based Theory of Natural Universal Computation. ArXiv: 2008.08814 [q-Bio].

Balsam, P.D., and Gallistel, C.R. (2009). Temporal maps and informativeness in associative learning. Trends Neurosci 32, 73–78.

Balsam, P.D., Drew, M.R., and Gallistel, C.R. (2010). Time and Associative Learning. Comp Cogn Behav Rev 5, 1–22.

Bangasser, D.A., Waxler, D.E., Santollo, J., and Shors, T.J. (2006). Trace Conditioning and the Hippocampus: The Importance of Contiguity. J Neurosci 26, 8702–8706.

Beylin, A.V., Gandhi, C.C., Wood, G.E., Talk, A.C., Matzel, L.D., and Shors, T.J. (2001). The role of the hippocampus in trace conditioning: temporal discontinuity or task difficulty? Neurobiol Learn Mem 76, 447–461.

Blanchard, T.C., Pearson, J.M., and Hayden, B.Y. (2013). Postreward delays and systematic biases in measures of animal temporal discounting. Proc. Natl. Acad. Sci. U.S.A. 110, 15491–15496.

Bradtke, S.J., and Duff, M.O. (1994). Reinforcement learning methods for continuous-time Markov decision problems. In Proceedings of the 7th International Conference on Neural Information Processing Systems, (Cambridge, MA, USA: MIT Press), pp. 393–400.

Brandon, S.E., Vogel, E.H., and Wagner, A.R. (2003). Stimulus representation in SOP: I. Theoretical rationalization and some implications. Behav Processes 62, 5–25.

Bright, I.M., Meister, M.L.R., Cruzado, N.A., Tiganj, Z., Buffalo, E.A., and Howard, M.W. (2020). A temporal record of the past with a spectrum of time constants in the monkey entorhinal cortex. PNAS 117, 20274–20283.

Carter, R.M., Hofstötter, C., Tsuchiya, N., and Koch, C. (2003). Working memory and fear conditioning. PNAS 100, 1399–1404.

Chittka, L., Geiger, K., and Kunze, J. (1995). The influences of landmarks on distance estimation of honey bees. Animal Behaviour 50, 23–31.

Clark, R.E., Manns, J.R., and Squire, L.R. (2002). Classical conditioning, awareness, and brain systems. Trends Cogn Sci 6, 524–531.

Coddington, L.T., and Dudman, J.T. (2018). The timing of action determines reward prediction signals in identified midbrain dopamine neurons. Nat. Neurosci. 21, 1563–1573.

Cohen, J.Y., Haesler, S., Vong, L., Lowell, B.B., and Uchida, N. (2012). Neuron-type-specific signals for reward and punishment in the ventral tegmental area. Nature 482, 85–88.

Daw, N.D., Courville, A.C., Tourtezky, D.S., and Touretzky, D.S. (2006). Representation and timing in theories of the dopamine system. Neural Comput 18, 1637–1677.

Dayan, P. (1993). Improving Generalization for Temporal Difference Learning: The Successor Representation. Neural Computation 5, 613–624.

De Corte, B.J., and Matell, M.S. (2016). Temporal averaging across multiple response options: insight into the mechanisms underlying integration. Anim Cogn 19, 329–342.

Desmond, J.E., and Moore, J.W. (1988). Adaptive timing in neural networks: The conditioned response. Biol. Cybern. 58, 405–415.

Doya, K. (2000). Reinforcement Learning in Continuous Time and Space. Neural Computation 12, 219–245.

Dylla, K.V., Galili, D.S., Szyszka, P., and Lüdke, A. (2013). Trace conditioning in insects-keep the trace! Front Physiol 4, 67.

Etscorn, F., and Stephens, R. (1973). Establishment of conditioned taste aversions with a 24-hour CS-US interval. Physiological Psychology 1, 251–259.

Frederick, S., Loewenstein, G., and O’Donoghue, T. (2002). Time Discounting and Time Preference: A Critical Review. Journal of Economic Literature 40, 351–401.

Gallistel, C.R., and Gibbon, J. (2000). Time, rate, and conditioning. Psychol Rev 107, 289–344.

Gallistel, C.R., Fairhurst, S., and Balsam, P. (2004). The learning curve: implications of a quantitative analysis. Proc Natl Acad Sci U S A 101, 13124–13131.

Gallistel, C.R., Craig, A.R., and Shahan, T.A. (2014). Temporal contingency. Behav Processes 101, 89–96.

Gallistel, C.R., Craig, A.R., and Shahan, T.A. (2019). Contingency, contiguity, and causality in conditioning: Applying information theory and Weber’s Law to the assignment of credit problem. Psychol Rev 126, 761–773.

George Ainslie (2001). Breakdown of Will (Cambridge: Cambridge University Press).

Gershman, S.J. (2018). The Successor Representation: Its Computational Logic and Neural Substrates. J. Neurosci. 38, 7193–7200.

Gershman, S.J., Moustafa, A.A., and Ludvig, E.A. (2014). Time representation in reinforcement learning models of the basal ganglia. Front. Comput. Neurosci. 7.

Gibbon, J., and Balsam, P. (1981). Spreading associations in time. In Autoshaping and Conditioning Theory, C.M. Locurto, H.S. Terrace, and J. Gibbon, eds. (New York: Academic), pp. 219–253.

Grossberg, S., and Schmajuk, N.A. (1989). Neural dynamics of adaptive timing and temporal discrimination during associative learning. Neural Networks 2, 79–102.

Hallam, S.C., Grahame, N.J., and Miller, R.R. (1992). Exploring the edges of Pavlovian contingency space: An assessment of contingency theory and its various metrics. Learning and Motivation 23, 225–249.

Hamid, A.A., Pettibone, J.R., Mabrouk, O.S., Hetrick, V.L., Schmidt, R., Vander Weele, C.M., Kennedy, R.T., Aragona, B.J., and Berke, J.D. (2015). Mesolimbic dopamine signals the value of work. Nat. Neurosci.

Hammond, L.J., and Paynter Jr, W.E. (1983). Probabilistic contingency theories of animal conditioning: A critical analysis. Learning and Motivation 14, 527–550.

Han, C.J., O’Tuathaigh, C.M., Trigt, L. van, Quinn, J.J., Fanselow, M.S., Mongeau, R., Koch, C., and Anderson, D.J. (2003). Trace but not delay fear conditioning requires attention and the anterior cingulate cortex. PNAS 100, 13087–13092.

Hinderliter, C.F., Andrews, A., and Misanin, J.R. (2012). The Influence of Prior Handling on the Effective CS-US Interval in Long-Trace Taste-Aversion Conditioning in Rats. Psychol Rec 62, 91–96.

Holland, P.C. (2000). Trial and intertrial durations in appetitive conditioning in rats. Animal Learning & Behavior 28, 121–135.

Kalmbach, A., Chun, E., Taylor, K., Gallistel, C.R., and Balsam, P.D. (2019). Time-scale-invariant information-theoretic contingencies in discrimination learning. Journal of Experimental Psychology: Animal Learning and Cognition 45, 280.

Kehoe, E.J., and Macrae, M. (2002). Fundamental behavioral methods and findings in classical conditioning. In A Neuroscientist’s Guide to Classical Conditioning, (Springer), pp. 171–231.

Kim, H.R., Malik, A.N., Mikhael, J.G., Bech, P., Tsutsui-Kimura, I., Sun, F., Zhang, Y., Li, Y., Watabe-Uchida, M., Gershman, S.J., et al. (2020). A Unified Framework for Dopamine Signals across Timescales. Cell 183, 1600–1616.e25.

Kobayashi, S., and Schultz, W. (2008). Influence of reward delays on responses of dopamine neurons. J. Neurosci. 28, 7837–7846.

Langille, J.J., and Gallistel, C.R. (2020). Locating the engram: Should we look for plastic synapses or information-storing molecules? Neurobiology of Learning and Memory 169, 107164.

Lattal, K.M. (1999). Trial and intertrial durations in Pavlovian conditioning: issues of learning and performance. J Exp Psychol Anim Behav Process 25, 433–450.

Ludvig, E.A., Sutton, R.S., and Kehoe, E.J. (2008). Stimulus representation and the timing of reward-prediction errors in models of the dopamine system. Neural Computation 20, 3034–3054.

Ludvig, E.A., Sutton, R.S., and Kehoe, E.J. (2012). Evaluating the TD model of classical conditioning. Learning & Behavior 40, 305–319.

Luzardo, A., Alonso, E., and Mondragón, E. (2017). A Rescorla-Wagner drift-diffusion model of conditioning and timing. PLOS Computational Biology 13, e1005796.

MacDonald, C.J., Lepage, K.Q., Eden, U.T., and Eichenbaum, H. (2011). Hippocampal “Time Cells” Bridge the Gap in Memory for Discontiguous Events. Neuron 71, 737–749.

Machado, A. (1997). Learning the temporal dynamics of behavior. Psychol Rev 104, 241–265.

Manns, J.R., Clark, R.E., and Squire, L.R. (2000). Awareness predicts the magnitude of single-cue trace eyeblink conditioning. Hippocampus 10, 181–186.

Matell, M.S., and Henning, A.M. (2013). Temporal memory averaging and post-encoding alterations in temporal expectation. Behav Processes 95, 31–39.

Matell, M.S., and Kurti, A.N. (2014). Reinforcement probability modulates temporal memory selection and integration processes. Acta Psychologica 147, 80–91.

Mello, G.B.M., Soares, S., and Paton, J.J. (2015). A Scalable Population Code for Time in the Striatum. Current Biology 25, 1113–1122.

Menzel, R. (2012). The honeybee as a model for understanding the basis of cognition. Nature Reviews Neuroscience 13, 758–768.

Miyazaki, K., Miyazaki, K.W., Sivori, G., Yamanaka, A., Tanaka, K.F., and Doya, K. (2020). Serotonergic projections to the orbitofrontal and medial prefrontal cortices differentially modulate waiting for future rewards. Sci Adv 6.

Mohebi, A., Pettibone, J.R., Hamid, A.A., Wong, J.-M.T., Vinson, L.T., Patriarchi, T., Tian, L., Kennedy, R.T., and Berke, J.D. (2019). Dissociable dopamine dynamics for learning and motivation. Nature 570, 65–70.

Momennejad, I., Russek, E.M., Cheong, J.H., Botvinick, M.M., Daw, N.D., and Gershman, S.J. (2017). The successor representation in human reinforcement learning. Nature Human Behaviour 1, 680–692.

Mondragón, E., Gray, J., Alonso, E., Bonardi, C., and Jennings, D.J. (2014). SSCC TD: A Serial and Simultaneous Configural-Cue Compound Stimuli Representation for Temporal Difference Learning. PLOS ONE 9, e102469.

Moore, J.W., Choi, J.-S., and Brunzell, D.H. (1998). Predictive timing under temporal uncertainty: the time derivative model of the conditioned response. Timing of Behavior: Neural, Psychological, and Computational Perspectives 3–34.

Morris, R.W., and Bouton, M.E. (2006). Effect of unconditioned stimulus magnitude on the emergence of conditioned responding. Journal of Experimental Psychology: Animal Behavior Processes 32, 371–385.

Namboodiri, V.M.K., Mihalas, S., and Shuler, M.G.H. (2014a). Rationalizing Decision-Making: Understanding the Cost and Perception of Time. Timing & Time Perception Reviews 1, 1–40.

Namboodiri, V.M.K., Mihalas, S., Marton, T.M., and Hussain Shuler, M.G. (2014b). A general theory of intertemporal decision-making and the perception of time. Front Behav Neurosci 8, 61.

Namboodiri, V.M.K., Mihalas, S., and Hussain Shuler, M.G. (2014c). A temporal basis for Weber’s law in value perception. Front Integr Neurosci 8, 79.

Namboodiri, V.M.K., Huertas, M.A., Monk, K.J., Shouval, H.Z., and Hussain Shuler, M.G. (2015). Visually cued action timing in the primary visual cortex. Neuron 86, 319–330.

Namboodiri, V.M.K., Mihalas, S., and Hussain Shuler, M.G. (2016a). Analytical Calculation of Errors in Time and Value Perception Due to a Subjective Time Accumulator: A Mechanistic Model and the Generation of Weber’s Law. Neural Comput 28, 89–117.

Namboodiri, V.M.K., Levy, J.M., Mihalas, S., Sims, D.W., and Hussain Shuler, M.G. (2016b). Rationalizing spatial exploration patterns of wild animals and humans through a temporal discounting framework. Proc. Natl. Acad. Sci. U.S.A. 113, 8747–8752.

Namboodiri, V.M.K., Otis, J.M., Heeswijk, K. van, Voets, E.S., Alghorazi, R.A., Rodriguez-Romaguera, J., Mihalas, S., and Stuber, G.D. (2019). Single-cell activity tracking reveals that orbitofrontal neurons acquire and maintain a long-term memory to guide behavioral adaptation. Nat. Neurosci. 22, 1110.

Narayanan, N.S., and Laubach, M. (2009). Delay activity in rodent frontal cortex during a simple reaction time task. J Neurophysiol 101, 2859–2871.

Niv, Y. (2009). Reinforcement learning in the brain. Journal of Mathematical Psychology 53, 139–154.

Pamir, E., Chakroborty, N.K., Stollhoff, N., Gehring, K.B., Antemann, V., Morgenstern, L., Felsenberg, J., Eisenhardt, D., Menzel, R., and Nawrot, M.P. (2011). Average group behavior does not represent individual behavior in classical conditioning of the honeybee. Learn. Mem. 18, 733–741.

Pamir, E., Szyszka, P., Scheiner, R., and Nawrot, M.P. (2014). Rapid learning dynamics in individual honeybees during classical conditioning. Front. Behav. Neurosci. 8.

Panoz-Brown, D., Corbin, H.E., Dalecki, S.J., Gentry, M., Brotheridge, S., Sluka, C.M., Wu, J.-E., and Crystal, J.D. (2016). Rats Remember Items in Context Using Episodic Memory. Current Biology 26, 2821–2826.

Panoz-Brown, D., Iyer, V., Carey, L.M., Sluka, C.M., Rajic, G., Kestenman, J., Gentry, M., Brotheridge, S., Somekh, I., Corbin, H.E., et al. (2018). Replay of Episodic Memories in the Rat. Current Biology 28, 1628–1634.e7.

Papachristos, E.B., and Gallistel, C.R. (2006). Autoshaped Head Poking in the Mouse: A Quantitative Analysis of the Learning Curve. Journal of the Experimental Analysis of Behavior 85, 293–308.

Pastalkova, E., Itskov, V., Amarasingham, A., and Buzsáki, G. (2008). Internally Generated Cell Assembly Sequences in the Rat Hippocampus. Science 321, 1322–1327.

Pavlov, I.P. (1927). Conditioned reflexes: an investigation of the physiological activity of the cerebral cortex (Oxford, England: Oxford Univ. Press).

Pearce, J.M., and Hall, G. (1980). A model for Pavlovian learning: variations in the effectiveness of conditioned but not of unconditioned stimuli. Psychol Rev 87, 532–552.

Petter, E.A., Gershman, S.J., and Meck, W.H. (2018). Integrating Models of Interval Timing and Reinforcement Learning. Trends in Cognitive Sciences 22, 911–922.

Rescorla, R.A. (1967). Pavlovian conditioning and its proper control procedures. Psychol Rev 74, 71–80.

Rescorla, R.A. (1968). Probability of shock in the presence and absence of cs in fear conditioning. Journal of Comparative and Physiological Psychology 66, 1–5.

Rescorla, R.A., and Wagner, A.R. (1972). A theory of Pavlovian conditioning: Variations in the effectiveness of reinforcement and nonreinforcement. Classical Conditioning II: Current Research and Theory 2, 64–99.

Russek, E.M., Momennejad, I., Botvinick, M.M., Gershman, S.J., and Daw, N.D. (2017). Predictive representations can link model-based reinforcement learning to model-free mechanisms. PLOS Computational Biology 13, e1005768.

Salz, D.M., Tiganj, Z., Khasnabish, S., Kohley, A., Sheehan, D., Howard, M.W., and Eichenbaum, H. (2016). Time Cells in Hippocampal Area CA3. J. Neurosci. 36, 7476–7484.

Schultz, W. (2016). Dopamine reward prediction error coding. Dialogues Clin Neurosci 18, 23–32.

Schultz, W., Dayan, P., and Montague, P.R. (1997). A Neural Substrate of Prediction and Reward. Science 275, 1593–1599.

Shankar, K.H., and Howard, M.W. (2012). A scale-invariant internal representation of time. Neural Comput 24, 134–193.

Stephens, D.W., and Krebs, J.R. (1986). Foraging Theory (Princeton University Press).

Sutton, R.S., and Barto, A.G. (1990). Time-derivative models of pavlovian reinforcement.

Sutton, R.S., and Barto, A.G. (1998). Introduction to Reinforcement Learning (Cambridge, MA, USA: MIT Press).

Takemoto, A., Miwa, M., Koba, R., Yamaguchi, C., Suzuki, H., and Nakamura, K. (2015). Individual variability in visual discrimination and reversal learning performance in common marmosets. Neuroscience Research 93, 136–143.

Thrailkill, E.A., Todd, T.P., and Bouton, M.E. (2020). Effects of conditioned stimulus (CS) duration, intertrial interval, and I/T ratio on appetitive Pavlovian conditioning. Journal of Experimental Psychology: Animal Learning and Cognition 46, 243–255.

Tiganj, Z., Jung, M.W., Kim, J., and Howard, M.W. (2017). Sequential Firing Codes for Time in Rodent Medial Prefrontal Cortex. Cerebral Cortex 27, 5663–5671.

Tiganj, Z., Cromer, J.A., Roy, J.E., Miller, E.K., and Howard, M.W. (2018). Compressed Timeline of Recent Experience in Monkey Lateral Prefrontal Cortex. Journal of Cognitive Neuroscience 30, 935–950.

Toure, M.W., Young, F.J., McMillan, W.O., and Montgomery, S.H. (2020). Heliconiini butterflies can learn time-dependent reward associations. Biology Letters 16, 20200424.

Tsao, A., Sugar, J., Lu, L., Wang, C., Knierim, J.J., Moser, M.-B., and Moser, E.I. (2018). Integrating time from experience in the lateral entorhinal cortex. Nature 561, 57–62.

Vogel, E.H., Brandon, S.E., and Wagner, A.R. (2003). Stimulus representation in SOP:: II. An application to inhibition of delay. Behavioural Processes 62, 27–48.

Wagner, A.R. (1981). SOP: A model of automatic memory processing in animal behavior. Information Processing in Animals: Memory Mechanisms 85, 5–47.

Ward, R.D., Gallistel, C.R., Jensen, G., Richards, V.L., Fairhurst, S., and Balsam, P.D. (2012). CS Informativeness Governs CS-US Associability. J Exp Psychol Anim Behav Process 38, 217–232.

Wystrach, A., Buehlmann, C., Schwarz, S., Cheng, K., and Graham, P. (2019a). Avoiding pitfalls: Trace conditioning and rapid aversive learning during route navigation in desert ants. BioRxiv 771204.

Wystrach, A., Schwarz, S., Graham, P., and Cheng, K. (2019b). Running paths to nowhere: repetition of routes shows how navigating ants modulate online the weights accorded to cues. Anim Cogn 22, 213–222.

Zhou, W., and Crystal, J.D. (2009). Evidence for remembering when events occurred in a rodent model of episodic memory. PNAS 106, 9525–9529.

